# Pediatric cerebrospinal fluid immune profiling distinguishes pediatric-onset multiple sclerosis from other pediatric-onset acute neurological disorders

**DOI:** 10.1101/2025.02.27.637541

**Authors:** Diego A. Espinoza, Tobias Zrzavy, Gautier Breville, Simon Thebault, Amaar Marefi, Ina Mexhitaj, Luana D. Yamashita, Mengyuan Kan, Micky Bacchus, Jessica Legaspi, Samantha Fernandez, Anna Melamed, Mallory Stubblebine, Yeseul Kim, Zachary Martinez, Caroline Diorio, Andreas Schulte-Mecklenbeck, Heinz Wiendl, Ayman Rezk, Rui Li, Sona Narula, Amy T. Waldman, Sarah E. Hopkins, Brenda Banwell, Amit Bar-Or

## Abstract

The cerebrospinal fluid (CSF) provides a unique glimpse into the central nervous system (CNS) compartment and offers insights into immune processes associated with both healthy immune surveillance as well as inflammatory disorders of the CNS. The latter include demyelinating disorders, such as multiple sclerosis (MS) and myelin oligodendrocyte glycoprotein antibody-associated disease (MOGAD), that warrant different therapeutic approaches yet are not always straightforward to distinguish on clinical and imaging grounds alone. Here, we establish a comprehensive phenotypic landscape of the pediatric CSF immune compartment across a range of non-inflammatory and inflammatory neurological disorders, with a focus on better elucidating CNS-associated immune mechanisms potentially involved in, and discriminating between, pediatric-onset MS (MS) and other pediatric-onset suspected neuroimmune disorders, including MOGAD. We find that CSF from pediatric patients with non-inflammatory neurological disorders is primarily composed of non-activated CD4+ T cells, with few if any B cells present. CSF from pediatric patients with acquired inflammatory demyelinating disorders is characterized by increased numbers of B cells compared to CSF of both patients with other inflammatory or non-inflammatory conditions. Certain features, including particular increased frequencies of antibody-secreting cells (ASCs) and decreased frequencies of CD14+ myeloid cells, distinguish MS from MOGAD and other acquired inflammatory demyelinating disorders.

## INTRODUCTION

In the non-inflammatory state, the central nervous system (CNS) contains immune cells involved in steady-state immune surveillance^1^, while in inflammatory disease states such as multiple sclerosis (MS), the abundance and profile of peripheral immune cells present within the CNS can vary and may include pathogenic immune cells ^1,2^. Cerebrospinal fluid (CSF), which is produced in the choroid plexus and drains into the subarachnoid granulations, is a fluid that envelops the brain in the subarachnoid space. It thus represents a CNS sub-compartment and may harbor cells that partially mirror disease processes occurring within the CNS tissue. To this end, prior studies have profiled CSF immune cells across neurological diseases including MS ^3–6^, Alzheimer’s disease^7^, and brain metastases^8^, among others (recently reviewed in ^9^), to elucidate disease-implicated immune mechanisms. However, while CSF cells from adult patients have been profiled extensively ^3–6,10–16^, there exists limited data on the CSF cellular compartment in the pediatric age group^5^, whether in health or across neurological disease states. The latter includes pediatric acquired inflammatory demyelinating syndromes (ADS), a collection of disorders in which immune responses are implicated in mediating CNS demyelination.

Pediatric-onset ADS include pediatric-onset MS (MS) which accounts for approximately 25% of all pediatric ADS diagnoses^17–23^; the more recently described MOGAD which accounts for approximately 30% of pediatric ADS diagnoses^17^; and individuals with ADS who do not meet MS, MOGAD, or neuromyelitis-optica spectrum disorder (NMOSD) diagnostic criteria (collectively referred to here as ‘other ADS’). MS, and particularly MOGAD, can share considerable overlap in their clinical presentation and disease course (with a relapsing course experienced in some patients with MOGAD). At the time of initial presentation, it may not be straightforward to distinguish the two conditions, and results of anti-MOG antibody testing that can be diagnostic for MOGAD may take days to weeks to become available. Like MS, patients with MOGAD can experience relapsing episodes of inflammatory demyelination. However, use of certain MS disease-modifying drugs may not be efficacious in prevention of MOGAD relapses and may even exacerbate its course^24–26^, pointing to distinct immunopathophysiological mechanisms underlying these two inflammatory demyelinating disorders, and the importance of distinguishing between them.

Here, we investigate the CSF immune cell compartment across a spectrum of pediatric-onset neurological disorders, including new-onset MS and MOGAD. We first characterize the immune compartment in non-inflammatory CNS conditions across the pediatric age-span. We then identify immune cell features distinguishing ADS from non-inflammatory and other (non-demyelinating) inflammatory disorders and further identify key cell-based immune features distinguishing MS from both MOGAD and other forms of ADS at the time of incident clinical presentation. In addition to establishing a foundational dataset describing the profile and dynamic nature of the cellular composition within non-inflammatory pediatric CSF throughout childhood and adolescence, we provide disease-specific insights into early CNS-associated inflammation in MS, MOGAD, other ADS and other neuroimmune disorders. We expect our findings to represent a useful resource for future studies into the pediatric CNS immune compartment in health and disease.

## RESULTS

### CSF immune cell profiling in the non-inflammatory pediatric CSF

We first established a standardized 16-parameter flow cytometric platform to profile immune cells from fresh pediatric CSF samples obtained at diagnostic lumbar punctures (**Fig. 1A**, see ***Methods***). This platform enabled us to identify and characterize major cell lineages (e.g. T cell, B cells, myeloid cells), as well as subsets within these lineages, from small volumes of pediatric CSF (0.5-7 mL) (**Supplementary Fig. 1**).

**Figure 1:**
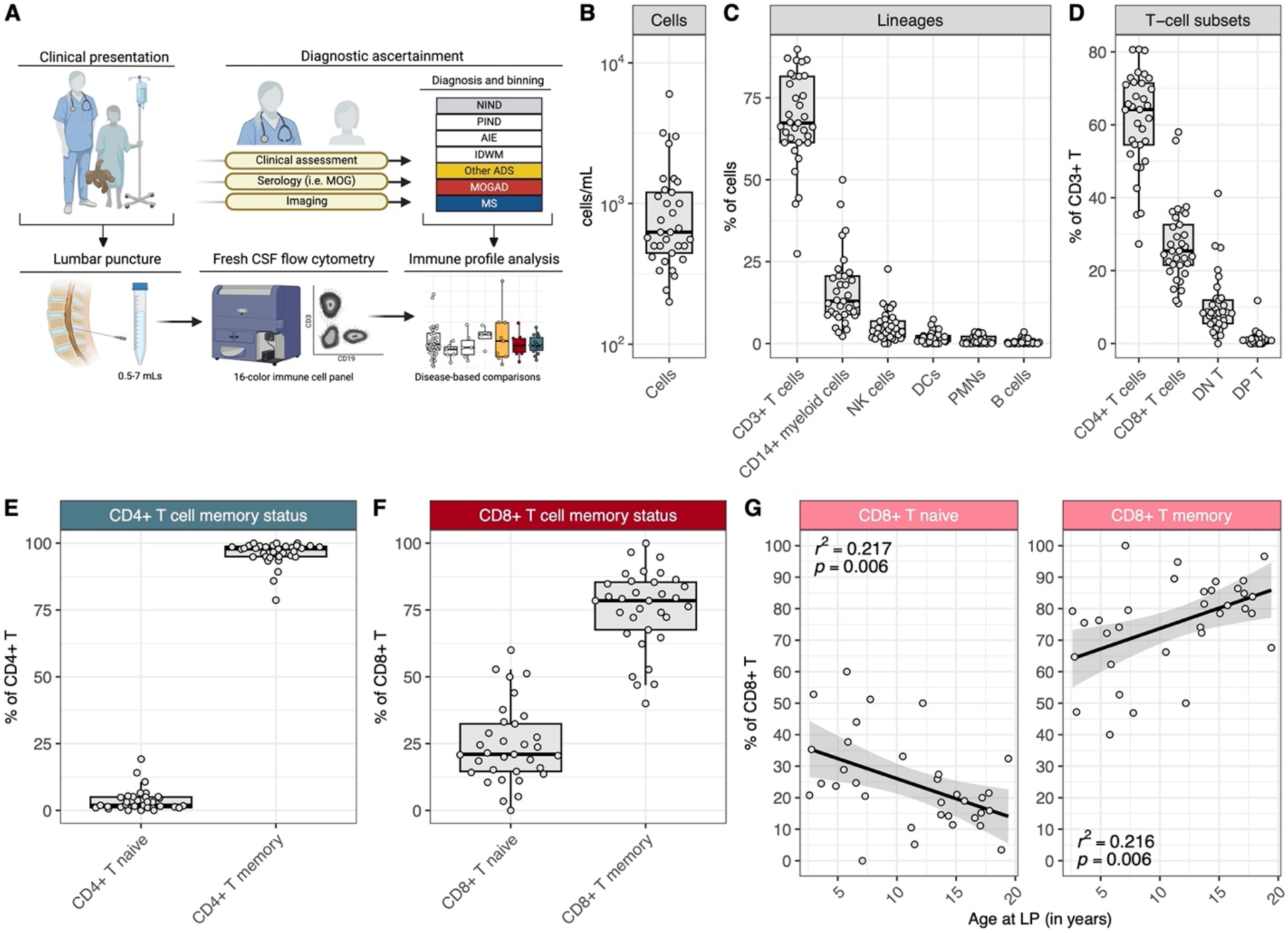
Pediatric NIND CSF is primarily composed of memory CD4+ and CD8+ T cells, with an age-associated increase in CD8+ memory T cell frequency. (**A**) Schematic of CSF sample procurement and flow cytometry data acquisition. (**B**) Cell count range (cells/mL) of NIND CSF (n=33). (**C**) Frequencies of CD3+ T cells, CD14+ myeloid cells, NK cells, DCs, PMNs, and B cells in NIND CSF. (D) Frequencies of T-cell subsets, as percent of CD3+ T cells, in NIND CSF. (**E**) Frequencies of CD4+ naive and memory T cells, as percent of CD4+ T cells, in NIND CSF. (**F**) Frequencies of CD8+ naive and memory T cells, as percent of CD8+ T cells, in NIND CSF and (**G**) across the NIND age span.

With this platform established, we next set out to determine the landscape of pediatric CSF across neurological conditions that are not considered primarily inflammatory in nature (i.e. non-inflammatory neurological disorders, or NIND, which include disorders such as primary headaches and idiopathic intracranial hypertension, among others (**Supplementary Table 1**). To this end, we analyzed the CSF flow cytometric profiles of 33 patients with subsequently ascertained diagnoses of NIND (**Table 1**). NIND CSF cell concentrations ranged from 0.2-6.0 cells/uL (median 0.63 cells/uL, **Fig. 1B**). The most abundant cell types within the cell fraction of the NIND CSF were CD3+ T cells (**Fig. 1C**, median 67.3%, CD45+/CD14-/CD19-/CD3+) and CD14+ myeloid cells (median 12.9%, CD45+/CD14+). Natural killer (NK cells 4.3%, CD45+/CD14-/CD3-/CD19-/CD56+), dendritic cells (DCs 1.6%, CD45+/CD14-/CD3-/CD19-/CD56-/HLA-DR+/CD11c+), B cells (0.6%, CD45+/CD14-/CD3-/CD19+) and polymorphonuclear cells (PMNs 0.3%, CD45+/SSChi, inclusive of neutrophils, eosinophils, and basophils) were detected at lower frequencies.

**Table 1.** Aggregated patient metadata by age and sex. NIND = non-inflammatory neurological disease, PIND = peripheral inflammatory neurological disease, AIE = autoimmune encephalitidies, IDWM = inherited disorders of white matter, other ADS = non-MS/non-MOGAD acquired demyelinating syndromes, MOGAD = myelin oligodendrocyte glycoprotein antibody-associated disease, MS = multiple sclerosis

| Category | Median Age | Age Range | Male Sex | Female Sex | Days from clinical episode onset to sampling (median for ADS groups only) | Days from clinical episode onset to sampling (range for ADS groups only) | Number in remission (MS only) |
| --- | --- | --- | --- | --- | --- | --- | --- |
| NIND | 12.2 | 2.6 - 19.4 | 21 | 12 | -- | -- | -- |
| PIND | 16.8 | 3.5 - 19.4 | 3 | 4 | -- | -- | -- |
| AIE | 14 | 8.7 - 19 | 2 | 4 | -- | -- | -- |
| IDWM | 10.1 | 1.3 - 17.2 | 4 | 0 | -- | -- | -- |
| other ADS | 9.3 | 0.7 - 17.9 | 8 | 2 | 4.5 | 1-40 | -- |
| MOGAD | 9.3 | 1.8 - 16.2 | 5 | 5 | 12 | 1-23 | -- |
| MS | 16.9 | 11.6 - 19.1 | 4 | 11 | 6 | 3-41 | 4 |

Within the CD3+ T-cell compartment of NIND CSF (**Fig. 1D**), most cells were CD4+ (median 67.2%), whereas CD8+ (median 23.6%) and double-negative (CD4-/CD8-; median 8.4%) T cells were less frequent. Double-positive (CD4+/CD8+) T cells were rarely observed (median 0.7%). Most CD4+ and CD8+ T cells were memory T cells (based on CD27 and CD45RA expression, median 97.8% and 77.0%, respectively, **Fig. 1E-F**), with larger variability in memory CD8+ T cell frequencies across donors. This variability appeared to be explained, at least in part, by a clear correlation between age and frequency of naive CD8+ T cells, such that youngest children with NIND had high frequencies of naive CD8+ T cells, with gradual increases in the frequency of memory CD8+ T cells observed with increasing age (**Fig. 1G**). Total CD8+ T-cell counts (cells/mL) remained stable over time (**Supplementary Fig. 2A**), indicating an age-associated shift in the balance of naive and memory CD8+ T cells in the non-inflamed pediatric CSF. We did not observe an age-associated pattern for naive and memory CD4+ T cells (**Supplementary Fig. 2B**).

Further assessment of the memory CD4+ and CD8+ T-cell compartments in patients with NIND revealed that memory CD4+ T cells were mostly CD27+/CD45RA-, consistent with a central-memory (CM) phenotype (**Supplementary Fig. 2C**) with the remainder being primarily effector-memory (EM) cells (CD27-/CD45RA-), and few CD4+ TEMRA cells (CD45RA+/CD27-) detected (**Supplementary Fig. 2C**). Most memory CD4+ T cells in NIND CSF were Th1-like (CXCR3+/CCR6-, **Supplementary Fig. 2D**). Few memory CD4+ T cells were activated (CD38+/HLA-DR+, **Supplementary Fig. 2E**). Similarly, most memory CD8+ T cells in the NIND patients had a CM phenotype (**Supplementary Fig. 2F**), and activated memory CD8+ T cells made up a minority of the memory CD8+ T-cell compartment (**Supplementary Fig. 2G**). Overall, pediatric NIND CSF was as expected of generally low cellularity, with most leukocytes being CD4+ T cells, most of which were memory T cells with a Th1-like phenotype and non-activated. CD8+ T cells were similarly mostly memory T cells and non-activated. We found that younger age correlated with an increased frequency of naive CD8+, but not CD4+, T cells.

### Elevations in B-cell counts highlight the increased cellularity of pediatric ADS CSF

Having established the CSF immune cell landscape in pediatric NIND, we next compared CSF cellular profiles of children with ADS (n = 33) to children with both NIND and a range of other inflammatory and non-inflammatory neurological disorders (**Table** 1, further detail in **Supplementary Table 1**). These disorders included peripheral inflammatory neurological disorders (PIND, n=7, which include inflammatory demyelinating disorders of the peripheral nervous system such as Guillain-Barre syndrome), autoimmune encephalitidies (AIE n = 6,, autoimmune inflammatory disorders of the CNS that are not necessarily demyelinating, such as NMDA-receptor encephalitis), and inherited disorders of white matter (IDWM, n = 4, genetic disorders resulting in abnormal growth or destruction of CNS white matter).

We first compared the absolute CSF cell counts of all children with ADS to children with NIND, PIND, AIE, and IDWM. We observed a statistically significant increase in the cellularity of ADS CSF compared to NIND, PIND, and AIE CSF (**Fig. 2A**). We next examined how individual lineages contributed to the increased cellularity in ADS CSF. We found that counts of total CD3+ T cells, CD14+ myeloid cells, NK cells, DCs, PMNs, and B cells showed statistically significant increases in ADS CSF when compared to NIND CSF (**Fig 2B**). Similarly, cell counts for these lineages tended to be increased when comparing ADS CSF to PIND, AIE, and IDWM CSF, though differences did not reach statistical significance for all comparisons (**Fig. 2B**). Of all examined lineages, the B-cell lineage exhibited the greatest relative enrichment in ADS CSF, with a greater than tenfold increase of B-cell counts in ADS CSF relative to B-cell counts in NIND, PIND, AIE, or IDWM CSF (**Fig. 2C**).

**Figure 2:**
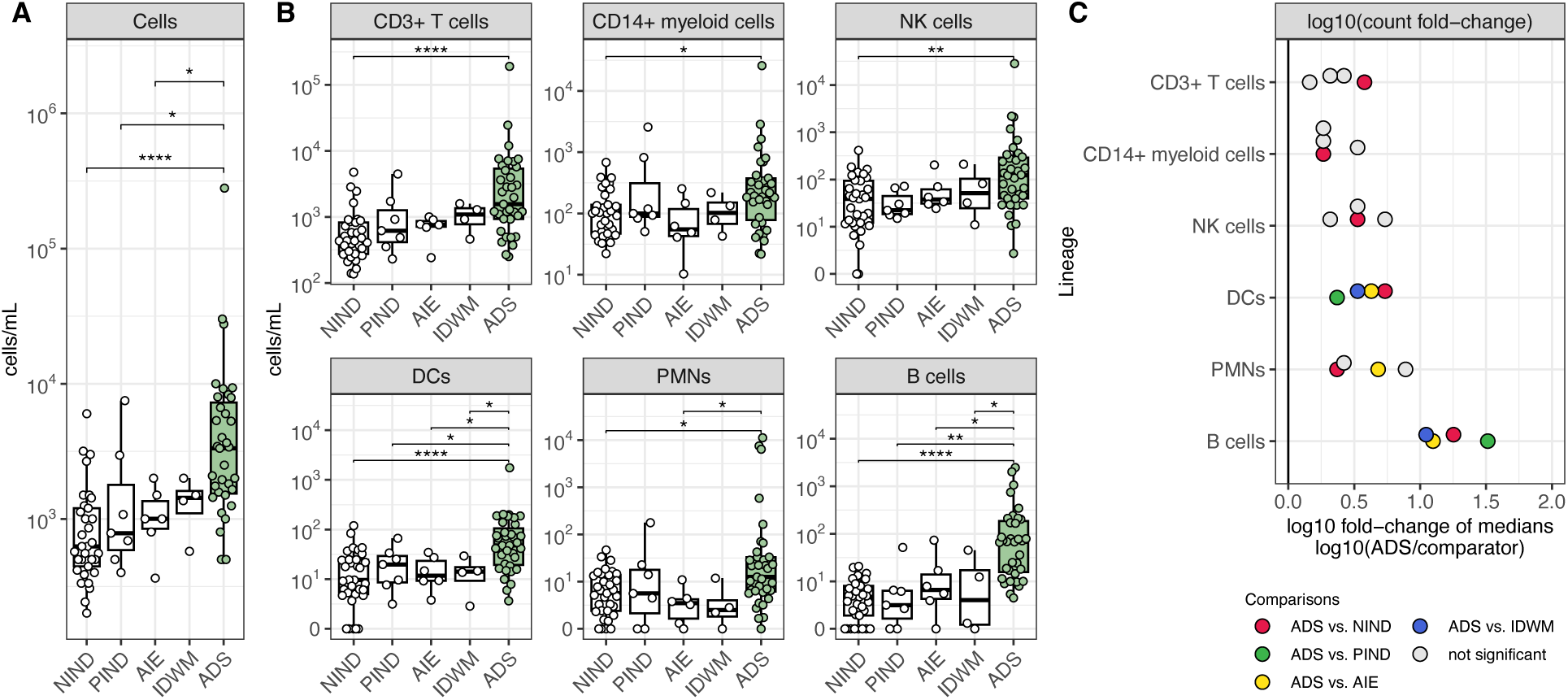
An increase in CSF B-cell counts best differentiates ADS from both other inflammatory and non-inflammatory neurological disease cohorts. **(A)** Cell counts (cells/mL) in NIND CSF (n=33), PIND CSF (n=7), AIE CSF (n=6), IDWM CSF (n=4), and ADS CSF (n=35). (**B**) Cell counts (cells/mL) for CD3+ T cells, CD14+ myeloid cells, NK cells, DCs, PMNs, and B cells across the same cohorts as **A**. (**C**) log10 of the fold change of median cell counts, comparing the median cell count in ADS to the median cell count in other cohorts for each lineage. For **Fig. 2A-B**, Wilcoxon-rank-sum test used to compare ADS to NIND, PIND, AIE, and IDWM independently (* = p < 0.05, ** = p < 0.01, *** = p < 0.001, **** = p < 0.0001). NIND = non-inflammatory neurological disease, PIND = peripheral inflammatory neurological disease, AIE = autoimmune encephalitidies, IDWM = inherited disorders of white matter, ADS = acquired demyelinating syndromes.

### MS CSF is distinguished from MOGAD and other ADS by a high frequency of antibody secreting cells and a decreased frequency of CD14+ myeloid cells

Having identified a pronounced B-cell enrichment in the CSF of children with ADS (**Fig. 2B-C**), we next assessed individual ADS diagnoses (MS, MOGAD, and other ADS). We observed that CSF B-cell counts were increased in each of MS, MOGAD, and other ADS CSF when independently compared to NIND CSF (**Supplementary Table 2**). Among these, B-cell frequencies were particularly elevated in MS CSF in comparison to MOGAD and other ADS CSF (**Fig. 3A**); B-cell absolute counts also appeared elevated in MS CSF in comparison to MOGAD and other ADS CSF, though this comparison did not reach statistical significance for MS vs. MOGAD, in part likely due to the large range of CSF B-cell and total cell counts observed in MOGAD patients (**Fig. 3A, Supplementary Fig. 3A**). We next sought to determine whether levels of antibody-secreting cells (ASCs), a B-cell subset that includes plasmablasts and plasma cells, differed between MS, MOGAD, and other ADS CSF. We found that both ASC frequencies and counts were increased in MS CSF when compared to MOGAD and other ADS (**Fig. 3B**). In fact, ASCs were frequently absent in MOGAD (absent in 5/10 samples) or other ADS CSF (absent in 5/10 samples) while almost always detected in MS CSF (**Fig. 3B**). Finally, frequencies and counts of B cells excluding ASCs (non-ASC B cells) appeared elevated in MS when compared to MOGAD and other ADS (**Supplementary Fig. 3B-C**). Together these findings point to elevated B-cell and particularly ASC levels as a feature distinguishing MS CSF not only from non-inflammatory disease controls but also from MOGAD and other ADS (**Fig. 3D**).

**Figure 3:**
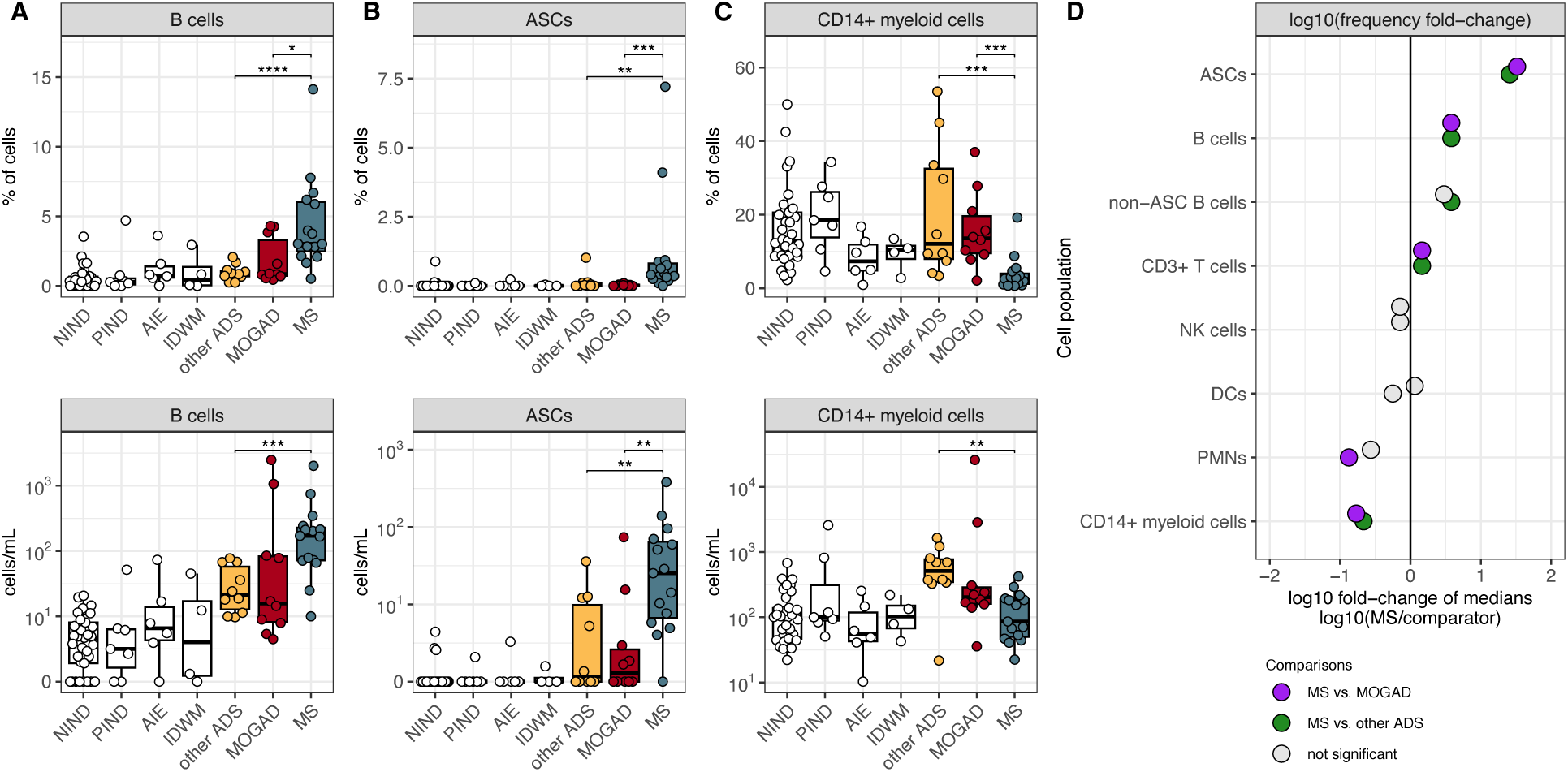
Pediatric MS CSF exhibits increased ASC frequencies and decreased CD14+ myeloid cell frequencies when compared to MOGAD and other ADS. **(A)** B-cell**, (B)** ASC, and (C) CD14+ myeloid cell frequencies and counts (cells/mL) in NIND CSF (n=33), PIND CSF (n=7), AIE CSF (n=6), IDWM CSF (n=4), other ADS CSF (n=10), MOGAD CSF (n=10), and MS CSF (n=15). (**D**) log10 of the fold change of median frequencies, comparing the median frequency in MS to the median frequency in other ADS and MOGAD for each cell population (lineage or subset). For **Fig. 3A-C**, Wilcoxon-rank-sum test used to compare MS to MOGAD, MS to other ADS, and MOGAD to other ADS (* = p < 0.05, ** = p < 0.01, *** = p < 0.001, **** = p < 0.0001). ASC = antibody secreting cell, NIND = non-inflammatory neurological disease, PIND = peripheral inflammatory neurological disease, AIE = autoimmune encephalitidies, IDWM = inherited disorders of white matter, other ADS = non-MS/non-MOGAD acquired demyelinating syndromes, MOGAD = myelin oligodendrocyte glycoprotein antibody-associated disease, MS = multiple sclerosis.

We further observed that frequencies of CSF CD14+ myeloid cells were lower in CSF of MS compared to both MOGAD and other ADS patients (**Fig. 3C-D**), which appeared to reflect elevations in CD14+ myeloid cell counts in the CSF of MOGAD and the other ADS patients, relative to NIND CSF (**Fig. 3C**, **Supplementary Table 2**).

CD3+ T-cell frequencies were enriched in MS CSF relative to MOGAD and other ADS CSF (**Supplementary Fig. 3B),** though no particular subset of T-cells clearly contributed this difference (**Supplementary Fig. 5A-B**). NK cell and DC frequencies and counts appeared to be similar across the three groups (**Supplementary Fig. 3B-C**). PMN frequencies appeared, at times, elevated in MOGAD and other ADS CSF, while largely absent in MS (**Supplementary Fig. 3B**). To explore this phenomenon further, we set a threshold (defined as >2 times the mean PMN frequency in NIND CSF) and assessed whether the presence of CSF PMNs above this threshold was associated with MOGAD/other ADS compared to MS (**Supplementary Fig. 3D**). Indeed, detection of CSF PMNs above this threshold was associated with a diagnosis of other ADS/MOGAD and essentially excluded MS (**Supplementary Fig. 3E**).

### Potential for the ASC to CD14+ myeloid cell ratio to distinguish MS CSF from MOGAD and other ADS CSF

Since an increase in ASC frequencies and a decrease in CD14+ myeloid cell frequencies appeared to best differentiate between CSF of MS versus MOGAD or other ADS patients (**Fig. 3D, Supplementary Fig. 4B**), we considered whether the ratio of ASC to CD14+ myeloid cell frequencies could be utilized to discriminate MS from MOGAD and other ADS, at the individual patient level. As expected, we found that the CSF ASC to CD14+ myeloid cell ratios (AMRs) were elevated in children with MS, both in comparison to MOGAD and to other ADS patients (p < 0.001, **Fig. 4A**). A previously reported ratio, termed the *neuroinflammatory composite score*^6^ (NCS), similarly incorporated several cellular measures, using both CSF (B-cell, CD14+ myeloid cell, CD56+ CD3+ NKT-cell frequencies and total cell counts) and blood (CD56^dim^ NK-cell frequencies) to distinguish between neuroinflammatory diagnoses (high composite scores), and non-inflammatory controls (lower composite scores). We utilized a CSF-only version of the NCS (coNCS, see Methods) and similarly found that this coNCS was also elevated in patients with MS, in comparison to MOGAD and other ADS (**Fig. 4B**). We found similar AMR and coNCS patterns distinguishing between patients who had received systemic glucocorticoid and/or intravenous immunoglobulin (IVIG) treatment prior to sampling versus those who had not (**Supplementary Fig. S5A-B**). We next assessed the performance of the AMR and the coNCS as classifiers for MS, when compared to either MOGAD, other ADS, or both by constructing receiver-operating characteristic (ROC) curves and found that both the AMR and coNCS performed well at discriminating MS from MOGAD and other ADS, with AUCs > 0.7 (**Fig. 4C**). The AMR, however, appeared superior to the coNCS in discriminating MS vs MOGAD (AUC=0.95, 0.80, respectively) and MS vs MOGAD/other ADS (AUC=0.94, 0.88, respectively). The AMR also provided excellent performance at distinguishing MS from other ADS (AUC=0.93), though the coNCS outperformed the AMR in this regard. (AUC=0.96). We were able to generate the full NCS in a subset of patients for whom we also had available blood CD56dim NK-cell frequencies and similarly observed that the full NCS was higher in MS relative to MOGAD and other ADS (**Supplementary Fig. 6A**) and overall performed similarly to the AMR (**Supplementary Fig. 6B**). Finally, we also evaluated the NCS companion “MS score”, which incorporates ASC presence and IgG synthesis rates. We found that it performed relatively poorly in our dataset (correctly classifying only 7 of 15 MS diagnoses; **Supplementary Table 3**). Overall, the two-parameter AMR we describe here provided excellent performance at discriminating MS CSF from MOGAD and other ADS CSF and performed at least as well as a previously reported neuro-immunological classifier reliant on additional parameters (NCS).

**Figure 4:**
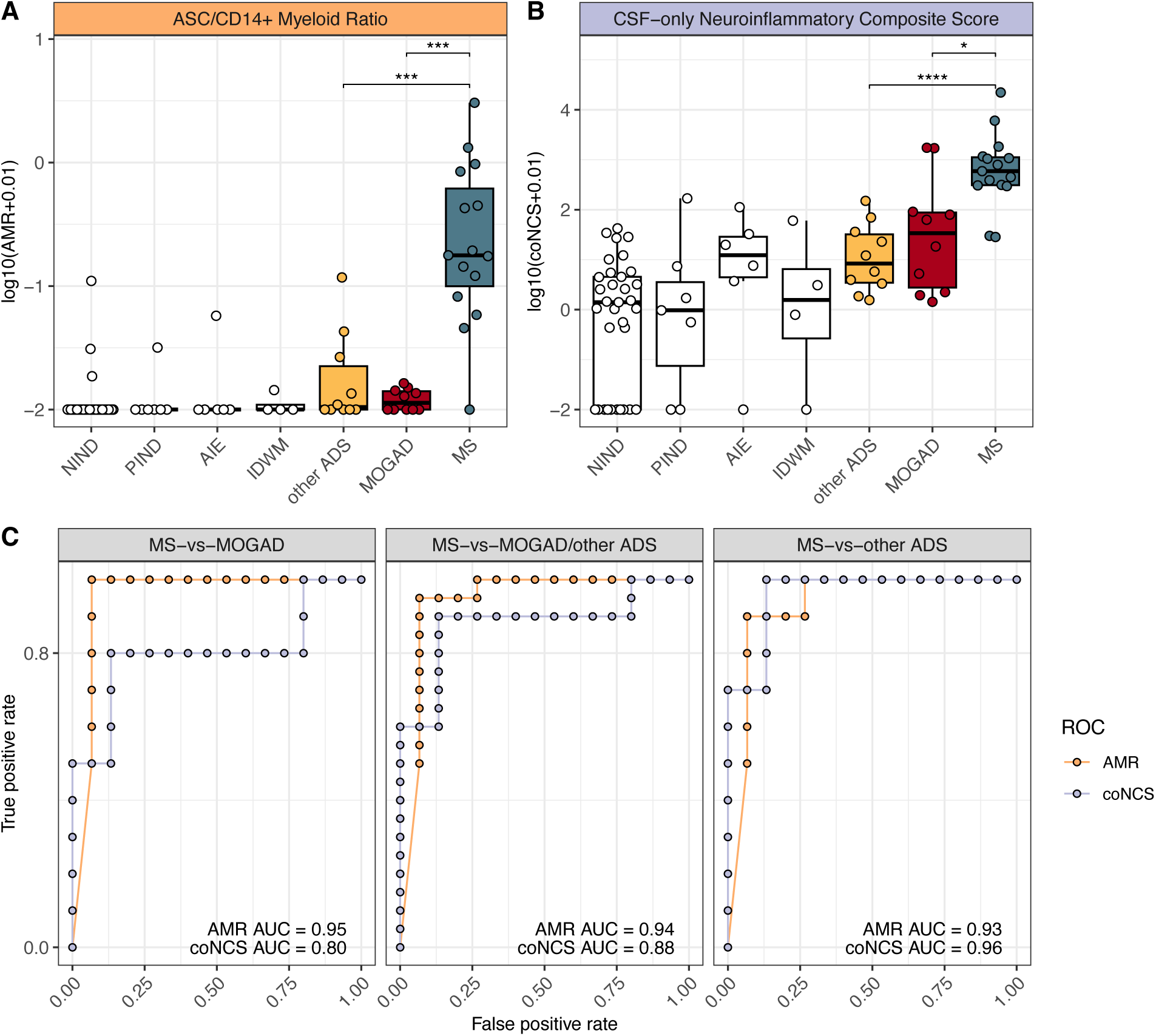
Ratio of CSF ASC to CSF CD14+ myeloid cell frequencies differentiates MS from MOGAD and other ADS. (**A**) Ratio of ASC frequencies to CD14+ myeloid cell frequencies (AMR) and (**B**) CSF-only neuroinflammatory composite scores (coNCS) across NIND CSF (n=33), PIND CSF (n=7), AIE CSF (n=6), IDWM CSF (n=4), and other ADS CSF (n=10), MOGAD CSF (n=10), and MS CSF (n=15). (**C**) Receiver operating characteristic (ROC) curves built for AMR or coNCS classifiers for the MS-vs-MOGAD/other ADS comparison, MS-vs-MOGAD comparison, and MS-vs-other ADS comparison, along with each corresponding area under the curve (AUC). For **Fig. 4A-B**, Wilcoxon-rank-sum used to compare MS to MOGAD, MS to other ADS, and MOGAD to other ADS (* = p < 0.05, ** = p < 0.01, *** = p < 0.001, **** = p < 0.0001). AMR = ASC to CD14+ myeloid cell ratio, coNCS = CSF-only neuroinflammatory composite score, NIND = non-inflammatory neurological disease, PIND = peripheral inflammatory neurological disease, AIE = autoimmune encephalitidies, IDWM = inherited disorders of white matter, other ADS = non-MS/non-MOGAD acquired demyelinating syndromes, MOGAD = myelin oligodendrocyte glycoprotein antibody-associated disease, MS = multiple sclerosis.

### Interactive exploration of pediatric CSF immune profiles

We next developed a public, web-based R-Shiny application for further analysis and visualization of pediatric CSF immune-profiles across a broader range of disease states (https://diegoalexespi.shinyapps.io/pedscsf/). By extending our CSF profiling to patients with viral encephalitis (VE) and patients with CNS-involved hemophagocytic lymphohistiocytosis (HLH), for example, we are able to observe a breadth of CD8+ T-cell activation states (**Supplementary Fig. S7**), such that patients with VE and HLH harbored elevated levels of CD8+ T-cell activation that are rarely observed in other disease states including other ADS, MOGAD, and MS (**Supplementary Fig. S7**). Such exploratory analyses can provide preliminary insight into further studies of the CSF immune compartment across disease states.

## DISCUSSION

Data on CSF cellular immune profiles in pediatric neurological disorders has been exceptionally limited. Here, we present a flow cytometric characterization of CSF immune cells across a spectrum of pediatric neurological disorders, identifying age-associated profiles in non-inflammatory neurological diseases that likely reflect steady-state immune surveillance, as well as disease-specific cellular signatures across a range of inflammatory states. We find that under normal, non-inflammatory conditions, pediatric CSF is not entirely acellular, with small numbers of immune cells predominantly composed of T cells, and with a CD8+ T-cell compartment that evolves with age. B cells are scarce or entirely absent in the CSF of non-inflammatory states while the CSF of children diagnosed with a range of ADS, exhibit notable increases in the number of B cells present. Among ADS presentations, a distinguishing feature of the CSF immune cell profile of children with MS compared to MOGAD and other ADS, is an increased frequency of ASCs and a decreased of frequency of CD14+ myeloid cells. Overall, these data represent the first characterization of pediatric CSF across both noninflammatory states and inflammatory neurological disorders and provides insight into normal immune surveillance as well as putative mechanisms of pediatric CNS disease.

Our characterization of the CSF profile across the pediatric age-span in non-inflammatory disorders provides us insight into age-associated patterns of CNS immune surveillance in the immunological steady-state. We find that most cells participating in CNS immune surveillance are CD4+ CM T cells across the pediatric age-span. These findings in the pediatric population are overall consistent with prior reports in adults that identified CM CD4+ and CD8+ T cells and CXCR3+ T cells as the primary cellular components of the NIND CSF immune compartment, with minimal to no B cells detected^27–29^. We extend these observations by finding that, across the pediatric age-span, CD8+ T cells shift from a naive (antigen-inexperienced) to memory (presumably antigen-experienced) phenotype with increasing age. This trend aligns with the observation that adult NIND CSF is largely devoid of naive CD8+ T cells^27^. Since the migration of immune cells from the periphery into the CNS is a regulated process and not a simple reflection of the periphery (as both CD4+ and CD8+ T cells are primarily naive throughout childhood^30^), this observation implies that naive CD8+ T cells earlier in life are endowed with a capacity to migrate into the CNS. One explanation is that these naive CD8+ T cells are “tissue-residency poised^31^” and mature into CD8+ tissue-resident memory T cells within the CNS that eventually saturate the CNS CD8+ tissue resident T-cell niche. Most parenchymal T cells in human brain (across health and disease states) appear to be CD8+ rather than CD4+, and express tissue residency markers^32^, consistent with the above phenomenon observed in CD8+ but not CD4+ T cells. It is also possible that what we observe as “naive” CD8+ T cells in the CSF, instead reflect a collection of antigen-experienced memory CD8+ T cells that share markers with naive CD8+ T cells^33,34^. In any event, it is evident that there exists an evolving recruitment of CD8+ T-cells of different phenotypes into the CNS across the pediatric age span, likely reflecting in part tandem immunological changes in CD8+ T-cell compartments beyond the CNS^35–37^.

We observe a significant elevation in CSF B-cell counts in children with ADS, beyond elevations observed for other lineages, and relative to both non-inflammatory and other inflammatory states. This elevation of B cells in ADS CSF could reflect a trafficking of peripheral B cells into the CNS where they may then participate in mechanisms associated with inflammatory demyelination; B cells are indeed present both in MS and MOGAD brain lesions^38,39^. The observation that B-cell numbers are elevated in both MS and MOGAD CSF does not necessarily mean they participate in these diseases through the same mechanism. In fact, B-cell depleting therapies, which are highly efficacious in preventing relapse activity in MS^40–42^, have had mixed results in preventing relapse activity in MOGAD^25,26^. Thus, it is plausible that these cells play distinct functional roles in different inflammatory demyelinating disease processes. Future studies could further delineate whether and how B cells in MS and MOGAD CSF differ with regards to, for example, their pro- and/or anti-inflammatory functions.

We found a particular enrichment of ASCs (the antibody-secreting B-cell subset) in MS CSF compared to other ADS and MOGAD. While ASC elevations have previously been described in MS CSF, prior studies have typically compared MS to non-inflammatory controls rather than to other inflammatory states^4,43–45^. Here, our direct comparisons with MOGAD and other ADS allowed us to demonstrate that increased presence of ASCs is a relatively distinct feature of MS CSF, rather than a feature of all ADS CSF. While peripheral ASCs may generate circulating antibodies to CNS proteins in certain ADS, such as the generation of circulating antibodies to MOG in MOGAD, the contribution of ASCs to disease in the MS CNS has not been fully elucidated and their function within the CNS may be distinct from that of peripheral ASCs. Of interest is whether and how the enrichment of ASCs in MS CSF may be associated somehow with presence (or possibly generation of) meningeal lymphoid aggregates, a feature also reported in the brains of patients with MS^46^ but not in MOGAD or other ADS. These aggregates, which are composed of T cells, B cells, plasma cells, and myeloid cells, are thought to contribute to chronic CNS-compartmentalized inflammation and progressive disease^47,48^, also a feature of MS, but not MOGAD^49^. If the presence of these ASCs in pediatric MS CSF were indeed associated with the presence or development of meningeal lymphoid aggregates, this could imply that the cellular machinery and anatomical structures associated with chronic CNS-compartmentalized inflammation in MS may already be developing very early in the MS disease spectrum.

We also identify decreased numbers of CD14+ myeloid cells in the CSF (and a concomitant increased ratio of ASCs to CD14+ myeloid cells) as a feature distinguishing MS from MOGAD and other ADS. Further work could dissect the pro/anti-inflammatory states of myeloid cells in the CSF of ADS patients including MOGAD, and assess their potential interactions with B-cell subsets, as such interactions are implicated in the context of CNS inflammatory disease^48^. Regardless of the processes that underlie the different CSF ratios of ASC to CD14+ myeloid cells, the ratio in our cohort distinguishes between MS and non-MS ADS including MOGAD and could be further investigated for its potential to provide an MS disease-specific CSF signature at time of initial CSF evaluation. The utility of this ratio may prove complementary to the previously developed CSF and blood “neuroinflammatory composite score (NCS)” that has been used in adults to distinguish between neuroinflammatory syndromes and non-inflammatory syndromes^6^.

We note that CSF CD4+ T cell subset profiles did not clearly distinguish between MS and other ADS or MOGAD. While prior reports in adults identified elevated frequencies of CD4+ regulatory T cells (Tregs) in MS CSF when compared to idiopathic intracranial hypertension (IIH) controls^4,50^, we did not find elevated Treg frequencies in MS CSF when compared to either NIND or to MOGAD or other ADS. Plausible explanations for this discrepancy include the more heterogenous cohort of NINDs included in our study and the differing ages of the patients across studies. While T cells observed in white matter lesions of MOGAD^39^ have been shown to be primarily CD4+ (in contrast to MS lesions, which are CD8+ T-cell dominant^38,51^), we did not find differences in CSF frequencies of CD4+ T cells or CD8+ T cells between MS and MOGAD patients.

While we were able to identify a CSF immune profile that distinguishes MS from MOGAD (and other ADS), we did not identify a unique CSF immune profile that clearly distinguished MOGAD patients. In prior studies^52,53^ PMNs (i.e. neutrophils or eosinophils) have been found to be elevated in MOGAD CSF. Here we identify that elevated frequencies of PMNs (while potentially exclusionary for diagnosis of MS), is a potential feature of both MOGAD and other ADS. The presence of PMNs in the CSF has been historically associated with anti-pathogen responses (typically to bacterial or also viral species) during CNS infection, yet the development of MOGAD or other ADS have not been firmly linked to a specific viral infection (including EBV^54^). It is plausible that the presence of PMNs in select MOGAD and other ADS CSF samples could hint at some anti-pathogen (potentially anti-viral) immune processes that, in parallel, also manifest as anti-self, inflammatory CNS demyelination processes.

Our study has several limitations. While we profiled children as proximal as possible to clinical disease onset, the underlying disease processes may already be well underway prior to clinical manifestations, which may be particularly true in MS where evidence for injury manifesting as elevated serum neurofilament light chain (NfL) levels is documented well prior to clinical disease onset^55^. Another limitation is absence of CSF profiles from children with neuromyelitis optica spectrum disorder (NMOSD), which is in the differential diagnosis of children with incident ADS presentations. Even though it is relatively uncommon in the pediatric age-range, NMOSD is thought to involve pathogenic antibodies, and it will be important to assess in the future whether CSF ASC levels are elevated or not and how the ASC to CD14+ myeloid cell ratio manifests in such children. Given that relapsing disease is a shared feature between NMOSD and MS (and some children with MOGAD), defining these profiles in NMOSD could provide additional insight into the pathophysiology of antibody-mediated ADS and disease chronicity. Finally, our cohort consists of children who underwent diagnostically-necessary lumbar punctures, which may not be wholly representative of all MS, MOGAD, or other ADS diagnoses. The diagnostic criteria for these disorders do not always necessitate CSF analyses, and the set of children who do undergo lumbar punctures may be enriched for those whose presentations elicit diagnostic uncertainty. Clear immune profile patterns nevertheless emerged in our data, such as those clearly separating MS from MOGAD and other ADS.

In sum, our findings provide insights into CSF immune cell profiles in the context of both normal immune surveillance and CNS non-inflammatory and inflammatory diseases, across the pediatric age-span. Our findings identify CSF cellular features that distinguish pediatric-onset MS from MOGAD and other ADS and may imply that some features of chronic inflammation are present early in the disease course. While no cellular features clearly delineated MOGAD from other ADS, the profiles observed in MOGAD and other ADS, as well as in MS, could reflect biological heterogeneity that may have implications for a patient’s clinical course. For example, a patient’s responsiveness to treatment(s) and risk of future relapse. Future studies could determine such “endophenotypes” by acquiring these CSF measures at the time of presentation and associating them with prospectively ascertained diagnoses, as has been done for peripheral blood profiles in adult MS^56^. As flow cytometric profiling of CSF samples becomes more widely available (driven by newly-developed CSF fixation protocols leveraged for multicenter studies^57,58^), CSF immune profiling has the potential to play an important complementary role in providing improved diagnosis and hence care of patients as part of a larger precision neuroimmunology approaches^59^.

## METHODS

### Patient eligibility and recruitment

Patients were recruited over a period of 3 years from inpatient and outpatient facilities at the Children’s Hospital of Philadelphia (CHOP). Patients were eligible and approached for enrollment (subject to staff availability) if they were undergoing a diagnostic lumbar puncture (LP) for a suspected neurological condition. Written informed consent was obtained from all donors. All protocols and consents were approved by the University of Pennsylvania Institutional Review Board.

### Acquisition, transport, and analysis of CSF samples

A total of 0.5-7 mLs of CSF per participant were obtained. CSF samples were collected in 15 mL polypropylene tubes at the time of LP, placed on ice and immediately transported to a single laboratory facility for prompt same-day processing and analysis. CSF from one HLH patient was collected from an external ventricular drain. Standardized operating procedures were consistent with consensus protocols for CSF analysis^60^.

### Flow cytometry of CSF cells

Upon arrival to the laboratory, 20 uL of CSF was placed on the Blood/Hemoglobin region of a Chemstrip (Chemstrip 10 SG, Cat. No. 11895362160) to evaluate for blood and/or hemoglobin contamination in the CSF sample. Samples with Chemstrip readings of ≥50 Erythrocytes/uL using either the blood or hemoglobin reading were not evaluated further. Evaluable samples were immediately spun at 4° C for 10 minutes at 400 RCF. CSF supernatant was then removed from the cell pellet, and the cell pellet was resuspended in 200 uL cold 1X PBS. 10 uL of sample was added to 10 uL of Methylene Blue and utilized for cell counting using a hemocytometer. Following cell counting, 800 uL of cold 1X PBS was added to the sample, and the sample was spun again at 4° C for 10 minutes at 400 RCF. Supernatant was discarded and the cell pellet was resuspended in 100 uL of cold 1X PBS for flow cytometry staining with a cocktail of 16 antibodies targeting CD45, CD14, CD3, CD19, CD56, HLA-DR, CD4, CD8, CD38, CD27, CD45RA, CD11c, CD127, CD25, CXCR3, and CCR6 (**Supplementary Table 4**) for 30 minutes at 4°C protected from light. Following the 30-minute stain, 900 uL of 1X cold PBS was added to the sample before another spin at 4° C for 10 minutes at 400 RCF. Supernatant was discarded and cell pellet was resuspended in 200-300 uL of 1X cold PBS and placed on ice prior to immediate analysis on an LSR Fortessa. Routinely, paired whole blood was stained and run on the Fortessa side-by-side with the CSF. BD FACSDiva software was utilized to collect flow cytometry data on the LSR Fortessa. Single-stain compensation tubes with beads were run prior to data collection and utilized for compensation on FlowJo software. After gating, frequencies were exported from FlowJo into .csv files for downstream statistical analysis.

### Statistical analysis

Statistical analysis was performed utilizing R software. When statistical comparisons were performed between conditions, a Wilcoxon-rank-sum test was utilized. The R packages *dplyr, ggplot2, magrittr, tibble, tidyr, ggpubr, pROC*, and *cowplot* were utilized for statistical analysis and figure generation.

### Neuroinflammatory composite score

A CSF-only neuroinflammatory composite score (coNCS, adapted from ^6^) was calculated with the following formula and used in **Fig. 4B**:

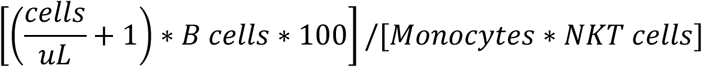

Where *cells/uL* = CSF cell concentration calculated from hemocytometer manual count, *B cells* = CSF B cell frequency as % of CSF lymphocytes, *Monocytes* = CSF CD14+ myeloid cell frequency as % of CSF cells, and *NKT cells* = CSF CD56+ CD3+ T-cell frequency as % of CSF lymphocytes. If either the denominator or numerator were 0, the coNCS was set to 0. For **Supplementary Fig. S6B**, the full NCS (with blood CD56dim NK) was utilized when blood CD56dim NK cell frequencies were available:

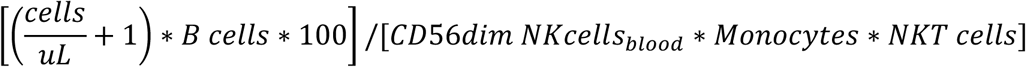

Where *CD56dim NK cells_blood_* = blood CD56dim NK cell frequency as % of blood lymphocytes and all other parameters identical to the coNCS. If either the denominator or numerator were 0, the full NCS was set to 0.

### Patient diagnoses and categorical classification

Patient diagnoses were ascertained in prospective follow-up by pediatric neurologists with extensive experience in pediatric neuroimmunology, and based on all available clinical and clinical laboratory data (but blinded to the CSF flow cytometry results). When possible, these diagnoses were then binned into five categories: non-inflammatory neurological disorders (NINDs), peripheral inflammatory neurological disorders (PIND), autoimmune encephalitidies (AIE), inherited disorders of white matter (IDWM). For acquired demyelinating syndromes (ADS), diagnoses were further characterized by presenting phenotype. The classification of specific diagnoses into categories is available within **Supplementary Table 1**. Within the ADS category, MOGAD was diagnosed utilizing the 2023 International MOGAD Panel proposed criteria^61^ and MS was diagnosed utilizing the 2017 McDonald criteria^62^. For children diagnosed with other ADS or MOGAD, CSF samples were determined to have been procured with a median of 4.5 days or 12 days, respectivley, from the onset of most recent episode of clinically active disease (**Supplementary Table 5**). For children diagnosed with MS, 4 of 15 were found to be in remission at the time of sampling, defined as the absence of clinical symptoms for at least 3 months prior to LP (**Supplementary Table 5**); for those with active clinical disease, CSF samples were determined to have been procured with a median of 6 days since the onset of most recent episode of clinically active disease (**Supplementary Table 5**). In **Supplementary Fig. S5**, other ADS, MOGAD, and CSF samples were further classified as *treated* if they had received any systemic glucocorticoid and/or IVIG treatment within the last 30 days prior to LP. Children with other ADS, MOGAD, or MS with any history of MS disease-modifying therapies were excluded from all analyses.

## DATA AVAILABILITY AND REPRODUCIBILITY

Complete R code (including R markdown notebooks with clear documentation) and final tables used to analyze and visualize our results have been deposited at https://github.com/diegoalexespi/espinozada_pediatricCSF2024.

## ACKNOWLEDGMENTS

This study was supported in part through the Melissa and Paul Anderson fund and Rosenblum donation (ABO, BB) with DAE supported in part by NIH Medical Scientist Training Program T32 GM07170, T32 G000046, F31AI167501, the Blavatnik Family Fellowship in Biomedical Research at the University of Pennsylvania, and the Penn Colton Center for Autoimmunity. TZ was supported in part by FWF Schrödinger grant (Nr: J4524). We would also like to thank the Children’s Hospital of Philadelphia Division of Child Neurology for their help in procuring CSF samples. Figure 1A was created with Biorender.com.

## AUTHOR CONTRIBUTIONS

DAE performed analysis. DAE, TZ, GB, ST, IM, LDY, YK, and ZM performed experiments. DAE, TZ, GB, ST, IM, MB, JL, SF, AnM, and MS assisted in sample transport and patient consenting. AmM ascertained patient diagnoses. DAE and MK designed the web-based R-shiny app for exploration of data. TZ, IM, CD, AS-M, HW, AR, and RL aided with analysis, protocol optimization, and panel development. ATW, SEH, and SN aided with patient recruitment and lumbar punctures. BB and ABO supervised the project. All authors contributed to the writing of the manuscript.

