## Supplementary Materials for "Pediatric cerebrospinal fluid immune profiling distinguishes pediatric-onset multiple sclerosis from other pediatric-onset acute neurological disorders"

### SUPPLEMENTARY MATERIALS: SUPPLEMENTARY FIGURES AND LEGENDS & SUPLEMENTARY TABLES

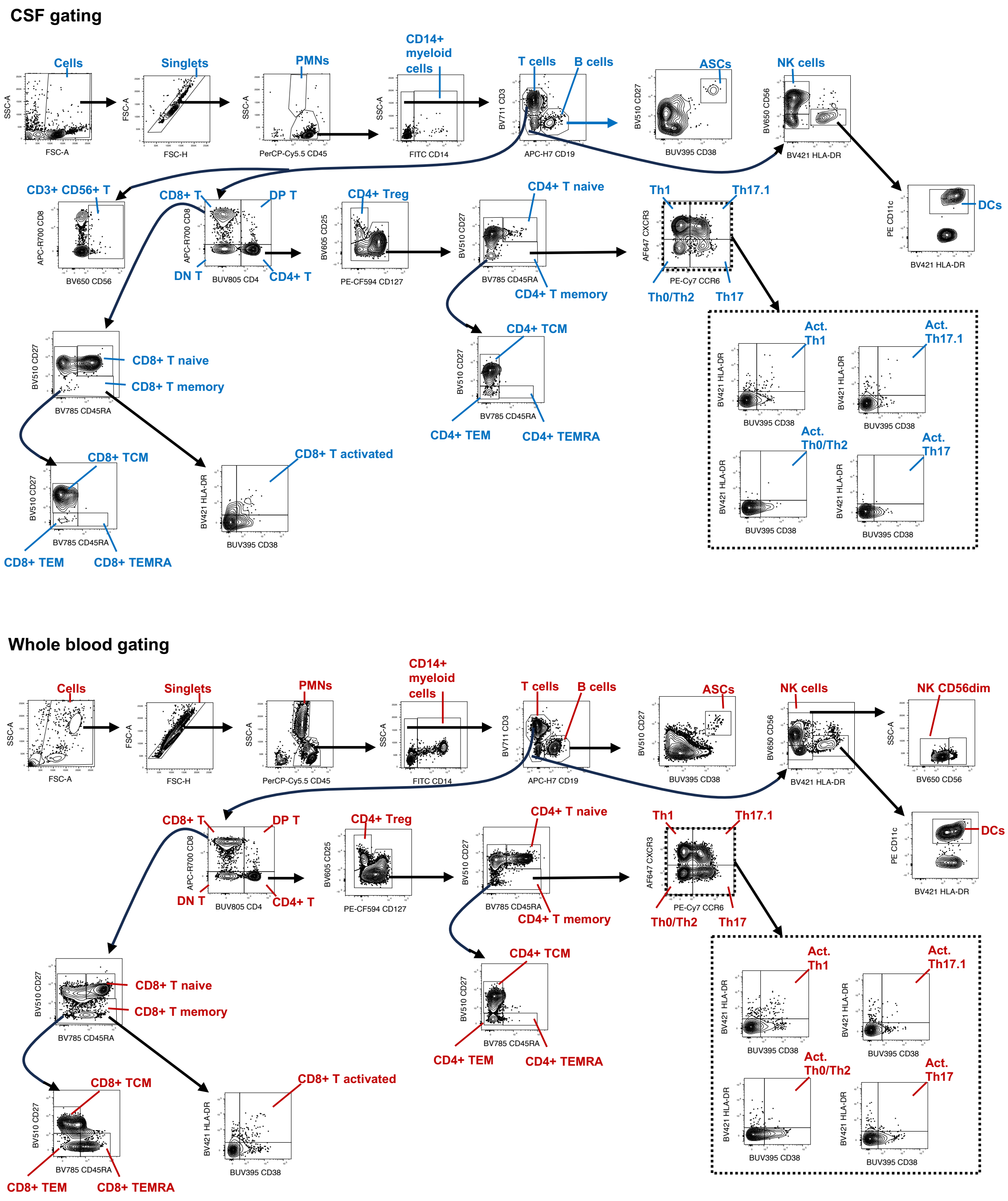


**Supplementary Figure 1: Gating strategy for 16-color flow cytometric platform.** Biaxial plots and gating strategy of flow cytometry results obtained using the 16-color flow cytometric platform for CSF and whole blood profiling in a representative example.  Act. = activated.

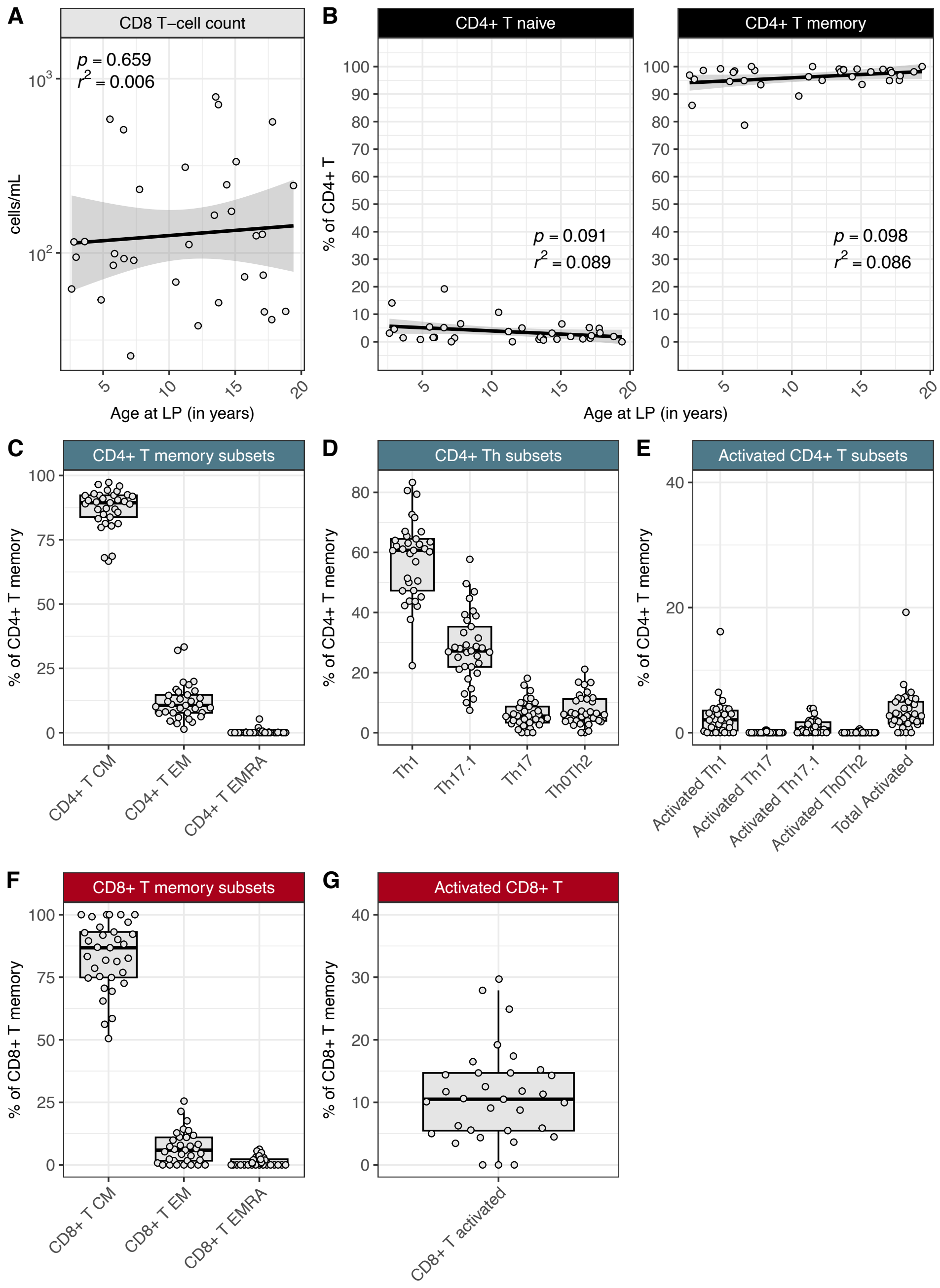


**Supplementary Figure 2: T-cell subsets in NIND CSF**. (**A**) Counts of total CD8+ T cells across the pediatric age-span in NIND CSF (n=33) with overlaid linear model. (**B**) Frequencies of naive and memory CD4+ T cells across the pediatric age-span in NIND CSF (n=33) with overlaid linear models. (**C**) Frequencies of central memory (CM), effector memory (EM), and effector memory CD45RA+ (EMRA) cells, as percent of memory CD4+ T-cells, in NIND CSF. (**D**) Distribution of

Th-status frequencies (based on CXCR3 and CCR6 expression) and their levels of activation based on HLA-DR/CD38 expression (**E**), as percent of memory CD4+ T-cells, in NIND CSF. (**F**) Frequencies of CM, EM, and EMRA cells as percent of memory CD8+ T-cells, in NIND CSF (n=33). (**G**) Frequencies of activated memory CD8+ T-cells in NIND CSF. NIND = non-inflammatory neurological disease.

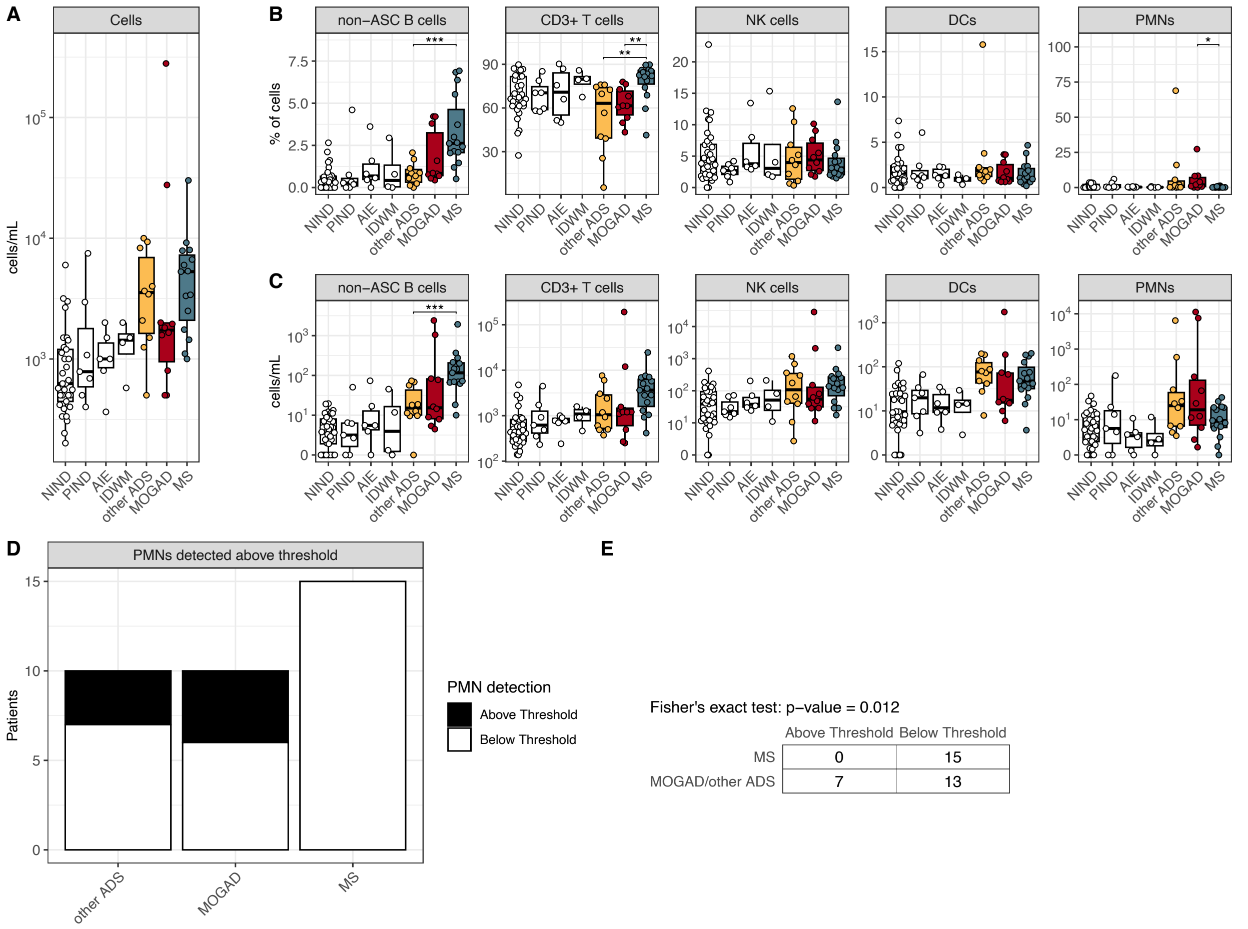


**Supplementary Figure 3: CSF cell counts and population frequencies across NIND, PIND, AIE, IDWM, other ADS, MOGAD, and MS.** (**A**) Cell counts (cells/mL) in NIND CSF (n=33), PIND CSF (n=7), AIE CSF (n=6), IDWM CSF (n=4), other ADS (n=10), MOGAD (n=10), and MS CSF (n=15). Frequencies (% of cells, **B**) and counts (cells/mL, **C**) of non-ASC B cells, CD3+ T cells, NK cells, DCs, PMNs across NIND, PIND, AIE, IDWM, other ADS, MOGAD, and MS CSF. (**D**) Frequency of PMN detection in patients above pre-defined threshold (see **Methods**) in other ADS, MOGAD, and MS CSF. (**E**) Fisher's exact test for the contingency table of PMN detection above pre-defined threshold, comparing MS CSF to MOGAD/other ADS CSF. For **Supplementary Fig. 3B-C**, Wilcoxon-rank-sum test used to compare MS to other ADS and MOGAD independently, and other ADS to MOGAD (* = p < 0.05, ** = p < 0.01, *** < 0.001,

**** < 0.0001). ASCs = antibody secreting cells., PMNs = polymorphonuclear cells NIND = non-inflammatory neurological disease, PIND = peripheral inflammatory neurological disease, AIE = autoimmune encephalitidies, IDWM = inherited disorders of white matter, other ADS = non-MS/non-MOGAD acquired demyelinating syndromes, MOGAD = myelin oligodendrocyte glycoprotein antibody-associated disease, MS = multiple sclerosis.

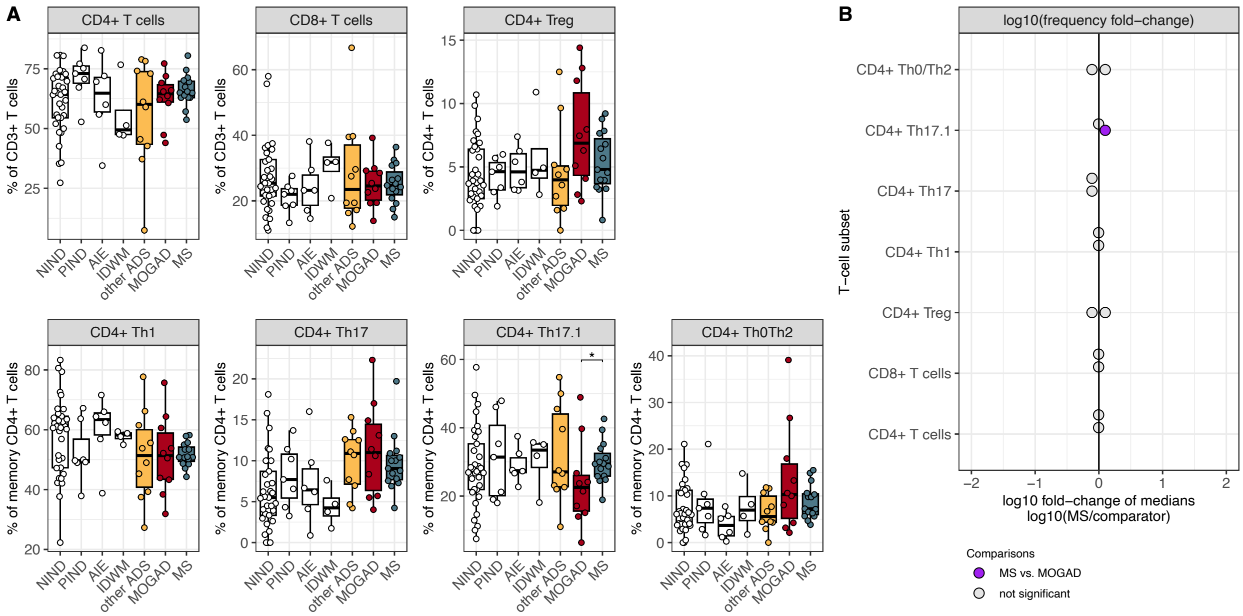


**Supplementary Figure 4: CSF T-cell subset frequencies across NIND, PIND, AIE, IDWM, other ADS, MOGAD, and MS.** (**A**) Frequencies of CD4+ T cells, CD8+ T cells, regulatory CD4+ T cells (CD4+ Treg), and CD4+ Th cell subsets in NIND CSF (n=33), PIND CSF (n=7), AIE CSF (n=6), IDWM CSF (n=4), other ADS (n=10), MOGAD (n=10), and MS CSF (n=15). (**B**) log10 of the fold change of median frequencies, comparing the median frequency in MS to the median frequency in other ADS and MOGAD for each T-cell subset. For **Supplementary Fig. 4A**, Wilcoxon-rank-sum test used to compare MS to other ADS and MOGAD independently, and other ADS to MOGAD (* = $p < 0.05$, ** = $p < 0.01$, *** < 0.001, **** < 0.0001). NIND = non-inflammatory neurological disease, PIND = peripheral inflammatory neurological disease, AIE = autoimmune encephalitidies, IDWM = inherited disorders of white matter, other ADS = non-MS/non-MOGAD acquired demyelinating syndromes, MOGAD = myelin oligodendrocyte glycoprotein antibody-associated disease, MS = multiple sclerosis.

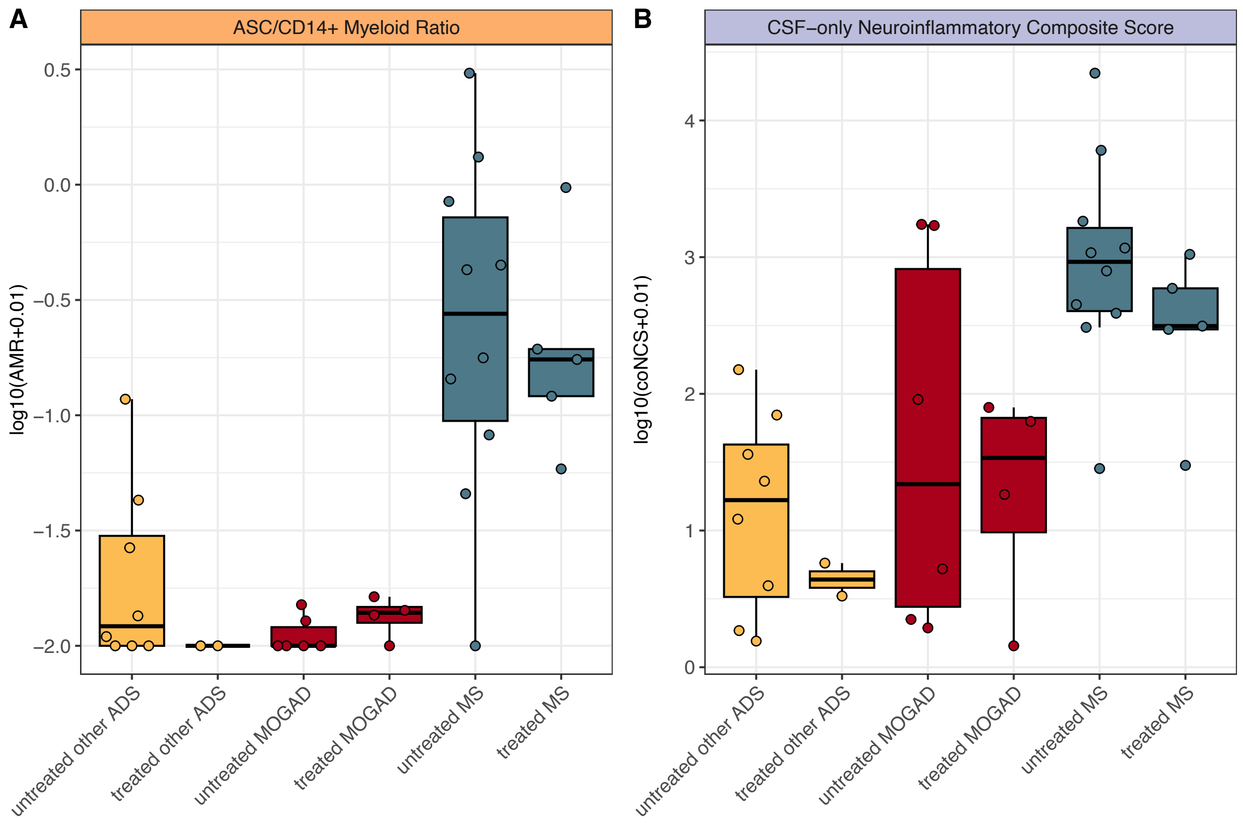


**Supplementary Figure 5: AMR and coNCS stratified by treatment status within ADS diagnoses.** (**A**) AMR and (**B**) coNCS across other ADS CSF (untreated n=8, treated n = 2), MOGAD CSF (untreated n=6, treated n=4), and MS CSF (untreated n=10, treated n=5). AMR = ASC to CD14+ myeloid cell ratio, coNCS = CSF-only neuroinflammatory composite score, other ADS = non-MS/non-MOGAD acquired demyelinating syndromes, MOGAD = myelin oligodendrocyte glycoprotein antibody-associated disease, MS = multiple sclerosis. “treated” indicates a CSF sample was drawn from a patient who had received any systemic glucocorticoid and/or IVIG treatment within 30 days prior to LP.

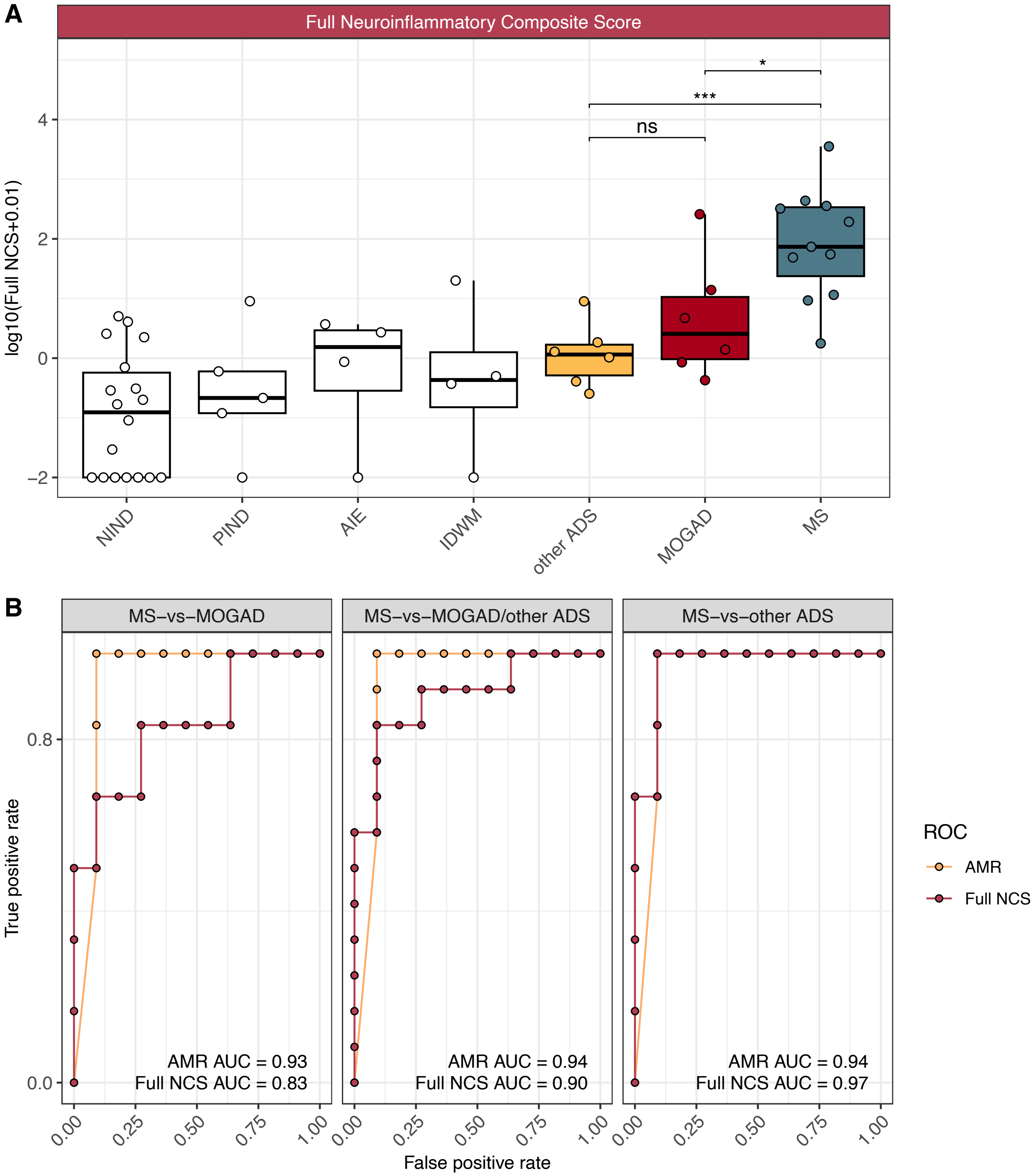


**Supplementary Figure 6: Evaluation of AMR compared to full NCS classifier.** (**A**) Values of the full NCS (utilizing blood measures, see **Methods**) across NIND CSF (n=18), PIND CSF (n=5), AIE CSF (n=4), IDWM CSF (n=4), and other ADS CSF (n=6), MOGAD CSF (n=6), and MS CSF (n=11). (**B**) Receiver operating characteristic (ROC) curves built for AMR or Full NCS classifiers for the MS-vs-MOGAD/other ADS comparison, MS-vs-MOGAD comparison, and MS-vs-other ADS comparison, along with each corresponding area under the curve (AUC). For **Supplementary Fig. 6A**, Wilcoxon-rank-sum used to compare MS to MOGAD, MS to other ADS, and MOGAD to other ADS (* = $p < 0.05$, ** = $p < 0.01$, *** = $p < 0.001$, **** = $p < 0.0001$). AMR = ASC to CD14+ myeloid cell ratio, NCS = neuroinflammatory composite score, NIND = non-inflammatory neurological disease, PIND = peripheral inflammatory neurological disease, AIE = autoimmune encephalitidies, IDWM = inherited disorders of white matter, other ADS = non-MS/non-MOGAD acquired demyelinating syndromes, MOGAD = myelin oligodendrocyte glycoprotein antibody-associated disease, MS = multiple sclerosis.

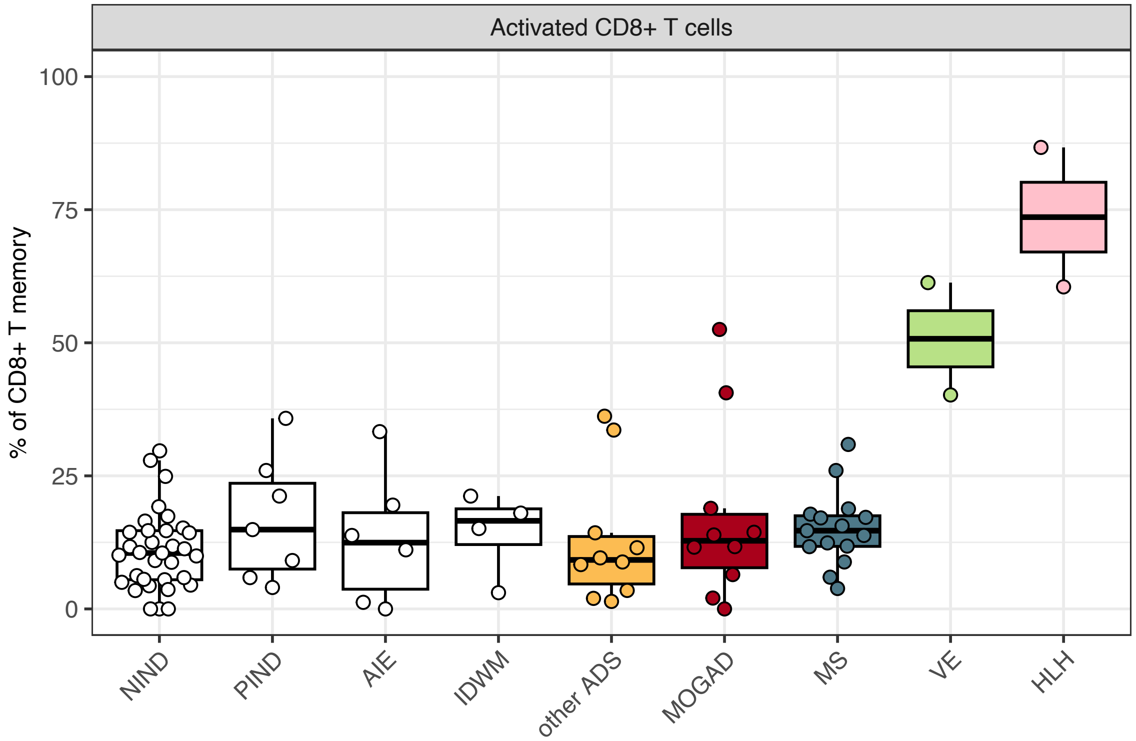


**Supplementary Figure 7: Elevated frequencies of CSF CD8+ T-cell activation in viral encephalitis and hemophagocytic lymphohistiocytosis.** CD8+ T-cell activation frequencies (as % of memory CD8+ T cells) in NIND CSF (n=33), PIND CSF (n=7), AIE CSF (n=6), IDWM CSF (n=4), other ADS (n=10), MOGAD (n=10), MS CSF (n=15), VE CSF (n=2), and HLH CSF (n=2). NIND = non-inflammatory neurological disease, PIND = peripheral inflammatory neurological disease, AIE = autoimmune encephalitidies, IDWM = inherited disorders of white matter, other ADS = non-MS/non-MOGAD acquired demyelinating syndromes, MOGAD = myelin oligodendrocyte glycoprotein antibody-associated disease, MS = multiple sclerosis, VE = viral encephalitis, HLH = hemophagocytic lymphohistiocytosis.

#### TABLE LEGENDS

**Supplementary Table 1. Patient diagnoses and classification.**

NIND = non-inflammatory neurological disease, PIND = peripheral inflammatory neurological disease, AIE = autoimmune encephalitidies, IDWM = inherited disorders of white matter, IIH = idiopathic intracranial hypertension, AIDP = acute inflammatory demyelinating polyneuropathy, CIDP = chronic inflammatory demyelinating polyneuropathy, GBS = Guillain-Barre syndrome, NMDARE = NMDA receptor encephalitis, HSV = herpes simplex virus, HLH = hemophagocytic lymphohistiocytosis, other ADS = non-MS/non-MOGAD acquired demyelinating syndromes, MOGAD = myelin oligodendrocyte glycoprotein antibody-associated disease, MS = multiple sclerosis, VE = viral encephalitis, HLH = hemophagocytic lymphohistiocytosis, ADEM = acute disseminated encephalomyelitis, ON = optic neuritis, TM = transverse myelitis, NOS = not otherwise specified.

**Supplementary Table 2. Wilcoxon rank-sum test results for B-cell and CD14+ myeloid cell comparisons between ADS groups and NIND.** NIND = non-inflammatory neurological disease, other ADS = non-MS/non-MOGAD acquired demyelinating syndromes, MOGAD = myelin oligodendrocyte glycoprotein antibody-associated disease, MS = multiple sclerosis.

**Supplementary Table 3. MS score calculated for pediatric samples.** MS score was cacluated as in Gross CC et al. and defined as plasma cells + intrathecal IgG synthesis, where: “plasma cells” = 1 if plasma cells are detected in CSF sample, 0 if not; “intrathecal IgG synthesis” = 1 if IgG synthesis rate elevated , 0 if not. Patient is classified as MS if their MS score is 2. IgG synthesis rate available for 9/10 other ADS patients, 7/10 MOGAD patients, and 15/15 MS patients. ASC presence in pediatric dataset used as equivalence to plasma cell presence in MS score. Other ADS = non-MS/non-MOGAD acquired demyelinating syndromes, MOGAD = myelin oligodendrocyte glycoprotein antibody-associated disease, MS = multiple sclerosis.

**Supplementary Table 4. 16-color flow cytometric panel targets, clones, fluorophores, dilution, and catalog numbers.**

**TABLES**

**Supplementary Table 1.**

| Category | Diagnosis |
|---|---|
| NIND | Altered mental status=4; Developmental delay=2; Developmental regression=2; Elevated optic nerves=2; Focal seizure=1; Headache=8; IIH=2; Landau Kleffner=1; Neuropathy=1; Papilledema=3; Pseudopapilledema=1; Psychiatric symptoms=6 |
| PIND | AIDP=1; Bell's palsy=1; Bilateral anterior uveitis=1; CIDP=1; CN VI palsy=1; GBS=1; Myasthenia gravis=1 |
| AIE | Antibody negative autoimmune encephalitis=3; NMDARE=3 |
| IDWM | Aicardi-Goutieres syndrome=1; HMBS-related leukoencephalopathy=1; Unknown genetic leukodystrophy=2 |
| ADS | Other ADS=10; MOGAD=10, MS=15 |
| VE | Eastern equine virus encephalitis=1; HSV encephalitis=1 |
| HLH | Isolated CNS HLH=1; XLP-HLH (due to SH2D1A mutation)=1 |

**Supplementary Table 2.**

| population_name | tested_value | group1 | group2 | p | p.label |
|---|---|---|---|---|---|
| B cells | cells/mL | NIND | other ADS | 0.00005 | **** |
| B cells | cells/mL | NIND | MOGAD | 0.00096 | *** |
| B cells | cells/mL | NIND | MS | 0.00000 | **** |
| CD14+ myeloid cells | cells/mL | NIND | other ADS | 0.00026 | *** |
| CD14+ myeloid cells | cells/mL | NIND | MOGAD | 0.01300 | * |
| CD14+ myeloid cells | cells/mL | NIND | MS | 0.84300 | ns |
| B cells | % of cells | NIND | other ADS | 0.01900 | * |
| B cells | % of cells | NIND | MOGAD | 0.00200 | ** |
| B cells | % of cells | NIND | MS | 0.00000 | **** |
| CD14+ myeloid cells | % of cells | NIND | other ADS | 0.89900 | ns |
| CD14+ myeloid cells | % of cells | NIND | MOGAD | 0.96600 | ns |
| CD14+ myeloid cells | % of cells | NIND | MS | 0.00000 | **** |

**Supplementary Table 3**

| Diagnosis | Patients with positive MS score | Total patients with available MS score |
|---|---|---|
| other ADS | 0 | 9 |
| MOGAD | 1 | 7 |
| MS | 7 | 15 |

**Supplementary Table 4**

| LASER NAME & FILTER | PARAMETER | TARGET | CLONE | DILUTION | CATALOG NUMBER |
|---|---|---|---|---|---|
| 530/30 Blue [B] | FITC | CD14 | MΦP9 | 1:20 | BD 347493 |
| 710/50 Blue [A] | PerCP-Cy5.5 | CD45 | HI30 | 1:50 | BD 564106 |
| 670/14 Red [C] | AF-647 | CXCR3 | G025H7 | 1:50 | BioLegend 353712 |
| 730/45 Red [B] | APC-R700 | CD8 | SK1 | 1:50 | BD 565192 |
| 780/60 Red [A] | APC-H7 | CD19 | SJ25-C1 | 1:50 | BD 560177 |
| 450/50 Violet [F] | BV421 | HLA-DR | G46-6 | 1:100 | BD 562804 |
| 525/50 Violet [E] | BV510 | CD27 | L128 | 1:50 | BD 563092 |
| 610/20 Violet [D] | BV605 | CD25 | 2A3 | 1:50 | BD 562660 |
| 660/20 Violet [C] | BV650 | CD56 | HCD56 | 1:50 | BioLegend 318344 |
| 710/50 Violet [B] | BV711 | CD3 | SK7 | 1:50 | BioLegend 344838 |
| 780/60 Violet [A] | BV786 | CD45RA | HI100 | 1:50 | BD 563870 |
| 379/28 UV [B] | BUV395 | CD38 | HB7 | 1:50 | BD 563811 |
| 820/60 UV [A] | BUV805 | CD4 | SK3 | 1:50 | BD 612888 |
| 586/15 YG [E] | PE | CD11c | Bu15 | 1:50 | BioLegend 337206 |
| 610/20 YG [D] | PE-CF594 | CD127 | HIL-7R-M21 | 1:50 | BD 562397 |
| 780/60 YG [A] | PE-Cy7 | CCR6 | 11A9 | 1:50 | BD 560620 |
